# Desynchronizing to be faster? Perceptual- and attentional-modulation of brain rhythms in sub-millisecond scale

**DOI:** 10.1101/329896

**Authors:** Yasuki Noguchi, Yi Xia, Ryusuke Kakigi

**Author notes:** Correspondence: Department of Psychology, Kobe University, 1-1 Rokkodai-cho, Nada, Kobe, 657-8501, Japan.

## Abstract

Neural oscillatory signals has been associated with many high-level functions (e.g. attention and working memory), because they reflect correlated behaviors of neural population that would facilitate the information transfer in the brain. On the other hand, a decreased power of oscillation (event-related desynchronization or ERD) has been associated with an irregular state in which many neurons behave in an uncorrelated manner. In contrast to this view, here we show that the human ERD is linked to the increased regularity of oscillatory signals. Using magnetoencephalography, we found that presenting a visual stimulus not only induced the ERD of alpha (8-13 Hz) and beta (13-30 Hz) rhythms in the contralateral visual cortex but also reduced the mean and variance of their inter-peak intervals (IPIs). This indicates that the suppressed alpha/beta rhythms became faster (reduced mean) and more regular (reduced variance) during visual stimulation. The same changes in IPIs, especially those of beta rhythm, were observed when subjects allocated their attention to a contralateral visual field. Those results revealed a new role of the ERD and further suggested that our brain regulates and accelerates a clock for neural computations by actively inducing the desynchronization in task-relevant regions.

## Introduction

Recent studies suggest that neural oscillatory signals could serve various important functions in the brain. Unit-cell recordings on animals showed that changes in amplitude and coherence of gamma-band oscillation are strongly linked to cognitive processes such as feature integration and attention (Engel et al., 2001; Fries, 2009), while oscillations in theta-band plays a key role in memory (Knyazev, 2007; Buzsaki and Moser, 2013). The close relationships between oscillatory signals and cognitive functions are also demonstrated in the human brain (Jensen et al., 2007; Palva and Palva, 2011). In electroencephalography (EEG) and magnetoencephalography (MEG), an increase in power of a given frequency band is called the event-related synchronization (ERS), which reflects cooperative or synchronized behaviors of a large number of neurons (Pfurtscheller and Lopes da Silva, 1999) that would facilitate the information transfer across neuronal groups (Fries, 2005). Consistent with this view, a spatial allocation of attention induced the gamma-band ERS in the sensory cortex of the contralateral hemisphere (Bauer et al., 2006), and stronger theta-band ERS in a memory encoding stage predicted better performances in later retrieval stage (Osipova et al., 2006).

In contrast to the ERS, a decrease in power of oscillatory signals is called the event-related desynchronization or ERD (Pfurtscheller and Lopes da Silva, 1999), being associated with uncorrelated (irregular) behaviors of neural population. Paradoxically, many studies have reported the ERD in brain regions that would play a central role in a given task (task-relevant regions). A well-known example is the alpha ERD over the occipital cortex in response to visual inputs (Berger, 1929). Recent studies showed that not only the alpha rhythm (8-12 Hz) but also the beta rhythm (13-30 Hz) showed prominent ERD to visual (Wyart and Tallon-Baudry, 2009; Minami et al., 2014; Kulashekhar et al., 2016), auditory (Cirelli et al., 2014; Fujioka et al., 2015; de Pesters et al., 2016), and somatosensory (Bauer et al., 2006; Fransen et al., 2016) stimuli. The alpha/beta ERD in the sensory areas was also induced by attention (van Ede et al., 2012). A voluntary allocation of attention to a visual hemifield reduced alpha/beta power in the contralateral visual cortex (Worden et al., 2000; Wyart and Tallon-Baudry, 2009; Bauer et al., 2014).

Why was the ERD, a neural signature of uncorrelated activities, observed in task-relevant regions? Previous studies resolved this issue by assuming an inhibitory role of alpha oscillation on neural processing (Jensen and Mazaheri, 2010; Klimesch, 2012). For example, Jansen and Mazaheri (2010) argued that alpha activity provides pulsed inhibition reducing the processing capabilities of a given area. A recent study also reported the inhibitory role of beta rhythm on neural processing (Shin et al., 2017). The alpha/beta ERD observed in task-relevant regions thus would reflect a release from this periodic (regular) inhibition, representing the active processing of sensory information in a contra-stimulus and contra-attention hemisphere.

Although this release-from-inhibition (disinhibition) hypothesis of alpha/beta ERD is widely accepted, here we explore a new role of ERD other than the disinhibition. Specifically, we focused on changes in *regularity* of neural rhythms caused by the ERD. Regularity (or periodicity) is one of the most fundamental information in oscillatory signals. Indeed, previous studies reported altered regularity of neural oscillations in patients with mental disorders such as Alzheimer’s disease (Gomez et al., 2007; Poza et al., 2012). We hypothesized that, if the alpha/beta ERD represents weakened (less-regulated) pulses of inhibition, this would be associated with decreased regularity of oscillations in the same frequency band.

Basic procedures of our MEG experiment are shown in **Figure 1**. Subjects performed the standard Posner task in which their attention was directed to either a left or right visual field (**Fig. 1A**). A central cue (arrow) and a peripheral task stimulus (Gabor patch) would induce alpha/beta ERD in a contralateral (contra-attention and contra-stimulus) hemisphere. To quantify the regularity, we measured inter-peak intervals (IPIs) of the oscillatory signals (**Fig. 1B**), depicting a distribution of their occurrences (**Fig. 1C**). For example, an IPI distribution of alpha rhythm (8-12 Hz) would be centered on 100 ms, because its central frequency is 10 Hz. When the alpha rhythm is highly regular, most IPIs would be located around the mean (100 ms), which results in a smaller variance of the distribution (**Fig. 1C**, top). In contrast, an irregular alpha wave would be indexed by a large variance because such irregular rhythm produces IPIs distant from 100 ms (**Fig. 1C**, bottom). One advantage of our method is that it can measure various types of changes in oscillatory signals simultaneously. For example, if the alpha/beta ERD causes an overall increase in neural processing speed, this would be detected as a decrease in a mean (rather than variance) of IPIs.

**Figure 1.**
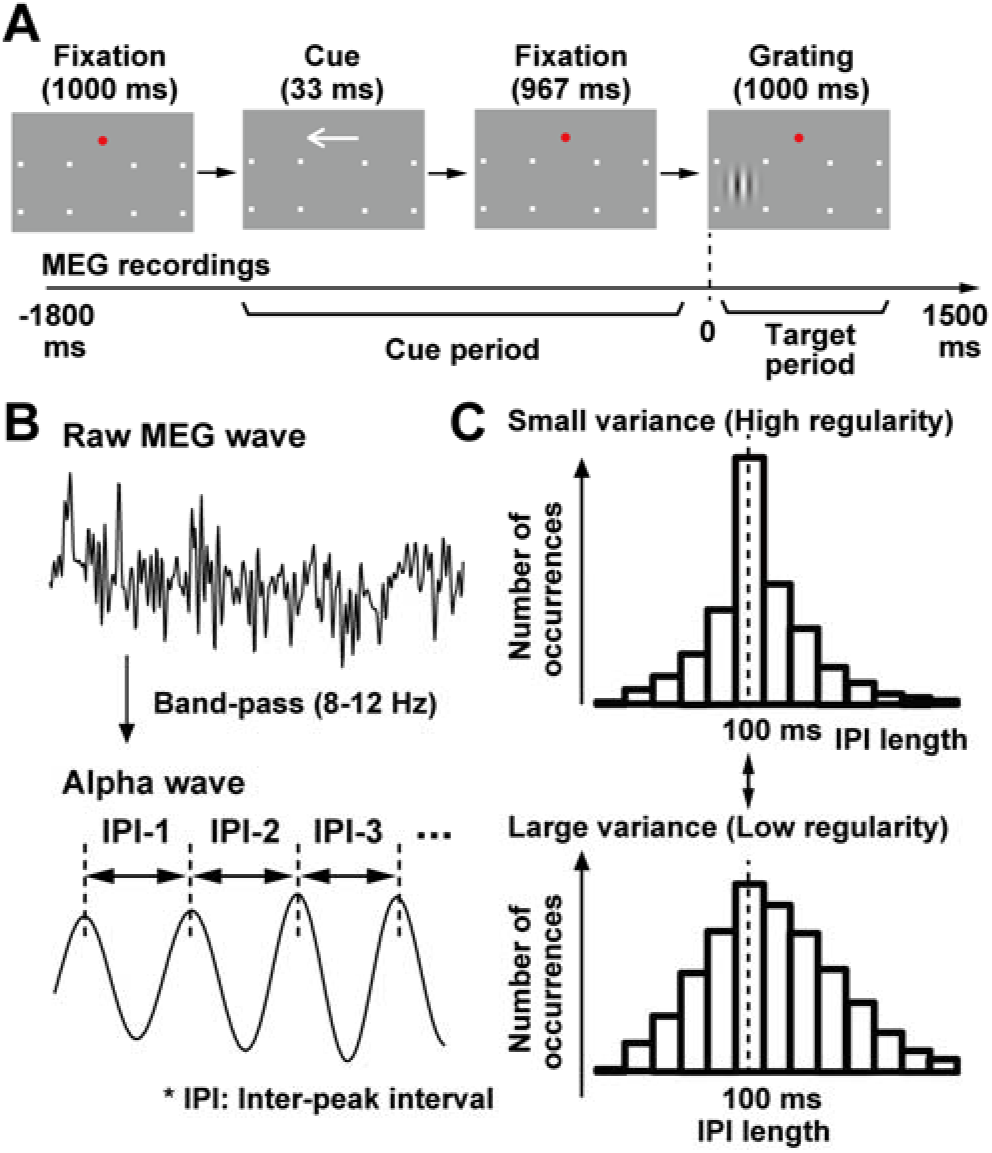
Experimental procedures. (**A**) Structures of one trial. A central cue (arrow) guided attention of subjects into either left visual field (LVF) or right visual field (RVF). Subjects pressed a button only when a grating with a target orientation (vertical or horizontal, variable across subjects) was presented at the attended hemifield. No response was required to a grating with a non-target orientation or a grating at the unattended hemifield. (**B**) Schematic illustrations of the inter-peak interval (IPI) analysis in the present study. First, oscillatory signals with a frequency band of interest (e.g. 8 - 12 Hz) were extracted with a band-pass filter. An IPI was defined as a time length between contiguous peaks of the filtered waveform. All IPIs within a period of interest (e.g. cue period) were pooled across trials. (**C**) Distributions of IPIs across all trials. The mean of the distribution should be around 100 ms (dotted line), because a central frequency of a pass band (8 - 12 Hz) is 10 Hz. When the oscillatory signals are highly regular, most IPIs would be located around the mean (upper panel). In contrast, an irregular alpha wave would produce many IPIs distant from the mean (lower panel). The irregularity of oscillatory signals can be thus indexed by a standard deviation (SD) of an IPI distribution.

## Materials and Methods

### Participants

Twenty-two human subjects (13 females, age: 20-54) participated in the present study. This sample size (N = 22) was larger than those in previous studies reporting alpha/beta ERD (N = 8-14) (Worden et al., 2000; Wyart and Tallon-Baudry, 2009; van Ede et al., 2012; Bauer et al., 2014). All had normal or corrected-to-normal vision. Informed consent was received from each participant after the nature of the study had been explained. All experiments were carried out in accordance with guidelines and regulations approved by the ethics committee of Kobe University, Japan.

### Stimuli and task

We used the Matlab Psychophysics Toolbox (Brainard, 1997; Pelli, 1997) to generate visual stimuli (refresh rate: 60 Hz) and to record button-press responses by participants. Each trial started with a screen of a red fixation point (0.19 ×0.19 deg) for 1000 ms (**Fig. 1A**). This fixation screen also had place holders (four white dots), one in the lower left visual field (LVF) and the other in the lower right visual field (RVF), indicating possible locations of an upcoming task stimulus (grating). To direct attention of participants, we then presented a cue stimulus (an arrow pointing leftward or rightward, 1.375 deg) over the central visual field for 33 ms, which was followed by another fixation screen for 967 ms. Finally, a grating with a vertical or horizontal orientation (size: 2 × 2 deg, center-to-fixation distance: 4.2 deg) was presented as a task stimulus either in LVF or RVF for 1000 ms. Participants were asked to move their attention covertly into a cued location, pressing a button as quickly as possible by their right hand when a grating with a target orientation (target grating) appeared there. The target orientation was vertical for a half of subjects (N = 11) and horizontal for the other half (N = 11). They were also instructed to hold the button-press response to a non-target grating in an attended hemifield and to totally ignore all gratings in an unattended hemifield.

A combination of a cued hemifield (left/right) and a position of the task stimulus (left/right) produced four types of trials: left cue & left stimulus (LL), left cue & right stimulus (LR), right cue & left stimulus (RL), right cue & right stimulus (RR). An experimental session comprised 72 trials in which those four types of trials (18 for each) were randomly intermixed. A ratio of target and non-target gratings was 3:15, meaning that participants should press the button for six times per session (three times for LL trials and three times for RR trials). Each participant underwent five sessions.

### MEG recordings

Neural activity was recorded with a whole-head MEG system (Vector-view, ELEKTA Neuromag, Helsinki, Finland). This system measured neuromagnetic signals from 102 positions over the scalp using 204 planer-type sensors (two sensors per recording position). One sensor measured latitudinal directions of changes in neuromagnetic signals while the other measured changes in longitudinal directions (sampling rate: 4 kHz, analogue band-pass filter: 0.1 – 330 Hz). Data from those two sensors at the same recording position were integrated in later analyses (see below). The MEG waveforms measured through those planar-type sensors represent neural activity in the cerebral cortex just below the recording position (Nishitani and Hari, 2002). Other details were shown in our previous publications (Suzuki et al., 2014; Noguchi et al., 2015).

The preprocessing of MEG data were performed with the Brainstorm (Tadel et al., 2011). First, neuromagnetic signals were segmented and classified into four conditions (LL, LR, RL, and RR). An epoch for the segmentation ranged from −1800 to 1500 ms relative to an onset of a task stimulus (**Fig. 1A**). Data in a pre-stimulus (post-cue) period from −1000 to 0 ms were used to investigate changes in oscillatory signals related to an allocation of attention, while changes related to visual perception were investigated in a post-stimulus (target) period from 0 to 1000 ms. The other periods at both ends (−1800 to −1000 ms and 1000 to 1500 ms) were used as a baseline or to avoid an edge effect of time-frequency transformation (see below). Trials in which a signal variation (difference between maximum and minimum values within a period of −1200 to 1000 ms) was larger than 4,500 fT/cm were excluded from further analyses. We also discarded trials in which a target grating was presented, because waveforms in those target trials contained noises resulting from manual button-press movements.

### Power analysis

We first analyzed changes in power of oscillatory signals over time, to confirm the attention- and perception-related alpha/beta ERD in previous studies (Worden et al., 2000; Wyart and Tallon-Baudry, 2009; Bauer et al., 2014; Minami et al., 2014; Kulashekhar et al., 2016). A time-frequency (TF) transform using complex Morlet wavelets was applied to the segmented MEG data of non-target trials (central frequency: 1 Hz, time resolution at full width at half maximum: 3 s), which converted an MEG waveform into a power spectrum of time (−1800 to 1500 ms) × frequency (1 to 100 Hz). Those TF spectra were then averaged across all trials and between latitudinal and longitudinal sensors at each recording position. To measure relative changes in power over time, we performed a baseline correction of the TF data. For each frequency, all data from −1500 to 1800 ms were converted into decibel (dB) change from a baseline, as shown by

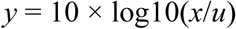

where *x* and *y* indicate powers before and after the baseline correction, and *u* indicates a mean power over a baseline period. The baseline period was set at −1200 to −1000 ms when we investigated power changes in the cue period (**Fig. 2**) and set at −200 to 0 ms when investigating changes in the target period (**Fig. 6**)

**Figure 2.**
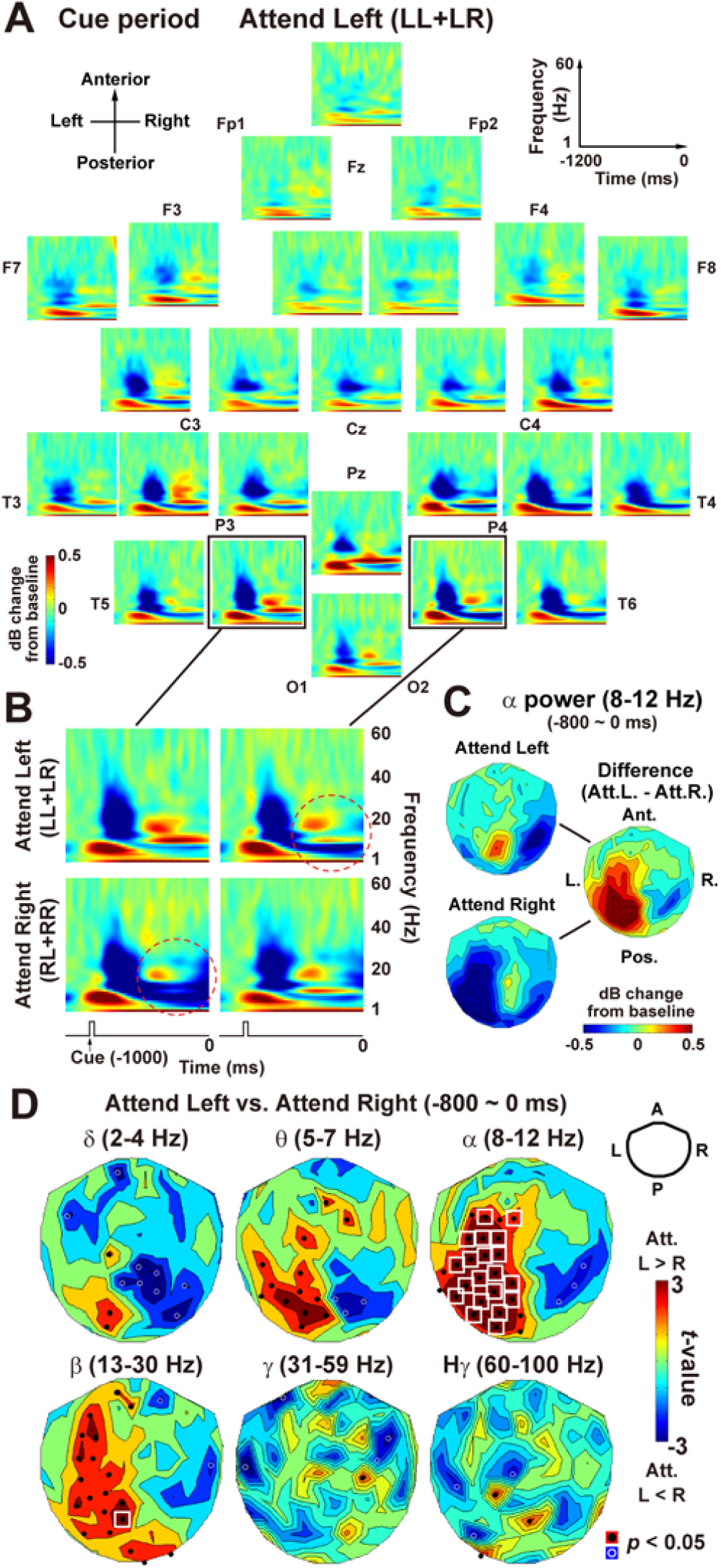
Results of the power analysis in the cue period. (**A**) Changes in power of oscillatory signals (1 to 60 Hz, vertical axis) over time (−1200 to 0 ms, horizontal axis) when subjects attended to LVF. Red and blue indicate an increase and decrease in power relative to a baseline period (−1200 to −1000 ms), respectively. We classified time-frequency (TF) spectra at 102 sensor positions into 26 groups (areas), with all spectra within the same group averaged together (areal-mean spectra). Approximate positions of the 26 areas are indicated by labels in the international 10-20 system (e.g. Cz). (**B**) Areal-mean spectra over the left and right visual regions in Attend-Left (upper panels) and Attend-Right (lower panels) conditions. Attending to LVF (RVF) suppressed alpha power over the right (left) visual cortex (dotted red circles). (**C**) Contour maps of alpha power over 102 sensor positions. A differential contour map (Attend-Left minus Attend-Right) was characterized by an increase (decrease) in alpha power in the left (right) visual cortex. (**D**) Statistical *t*-maps over 102 sensor positions. For each of six frequency bands from delta to high-gamma, oscillation powers are compared between Attend-Right and Attend-Left conditions. Black and white dots indicate sensors showing a significant (*p* < 0.05) difference. White rectangles denote sensors with a significant difference after a correction of multiple comparisons. An event-related desynchronization (ERD) in the contra-attention hemisphere was clearly observed in alpha band (upper right). Similar patterns were also seen in delta, theta, and beta bands.

### Source estimation

We estimated anatomical source locations of alpha suppression (**Fig. 6D**) with the minimal norm (MN) approach in Brainstorm. First, a spherical head model of each subject was constructed, using positional information of MEG sensors and a template brain of the Montreal Neurological Institute. This model for forward solutions was then inverted with the MN approach, which converted neuromagnetic waveforms at 204 sensors into a 4D current density map (cortical activation map) of multiple dipoles placed over the cortical surface. Finally, those changes in the current density over time were transformed into power spectrum with the Welch’s method (power density map). Noise covariance matrix for those MN estimations was computed using MEG signals in a pre-cue baseline period (−1200 to −1000 ms).

For second-level (group-level) analyses across 22 subjects, the power density map of each subject was exported to Statistic Parametric Mapping (SPM) 12. Brain regions showing the alpha suppression to a contralateral task stimulus were estimated by the random-effect analysis (voxel-wise *t*-tests) comparing Stimulus-Left condition (average of LL and RL trials, expressed as “LL+RL” hereafter) and Stimulus-Right condition (“LR+RR”).

### Inter-peak interval analysis

We evaluated regularity of neuromagnetic waveforms using the inter-peak interval (IPI) analysis. First, a frequency band of interest was defined based on results of the power analysis. For example, we found ERD in the contra-attention hemisphere in delta (2-4 Hz), theta (5-7 Hz), alpha (8-12 Hz), and beta (13-30 Hz) bands (**Fig. 2D**). The frequency band of interest was thus defined as 2-30 Hz (**Fig. 3**, left panels), to investigate how those ERDs affected oscillatory rhythms in the same band. As shown in **Figure 1B**, we applied a band-pass filter (Butterworth, zero-phase) of 2-30 Hz to raw MEG waveforms, identifying peaks and their intervals (IPIs). All IPIs within a period of interest (e.g. cue period) were pooled across trials and between latitudinal and longitudinal sensors at the same recording position. We then analyzed four parameters of an IPI distribution (**Fig. 1C**); mean, standard deviation (SD), median, and coefficient of variation (CV). The CV is the SD divided by the mean IPI, which represents the irregularity (variance) of neural signals normalized to a mean (Taube, 2010). A higher CV indicates lower regularity of signals.

**Figure 3.**
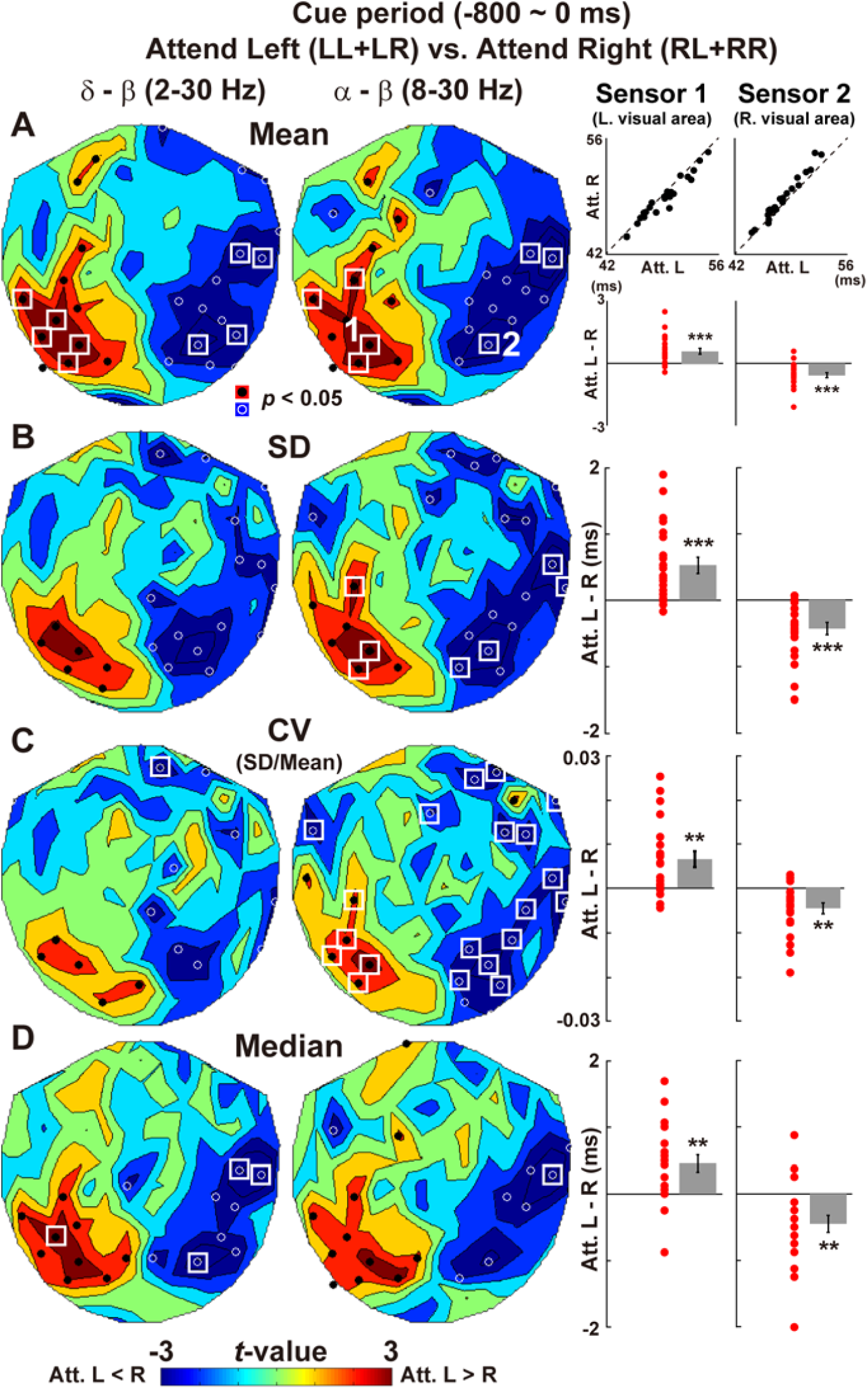
Results of the IPI analysis in the cue period. Sensor-space *t*-maps (Attend L vs. R) are shown for four parameters of an IPI distribution; mean (**A**), SD (**B**), coefficient of variation (**C**), and median (**D**). The coefficient of variation (CV) is the SD divided by the mean IPI, which represents the irregularity of neural signals (Taube, 2010). Left *t*-maps showed results when IPIs were measured using a band-pass filter of 2-30 Hz (delta to beta), while right *t*-maps showed results of IPIs measured through a filter of 8-30 Hz (alpha to beta). Data of individual subjects at representative sensors are shown in right panels (Sensor 1 over the left visual area and Sensor 2 over the right visual area, see the right *t*-map in **A** for their locations). For mean IPI (**A**), individual data (N = 22) in Attend-Left (*x*) and Attend-Right (*y*) conditions are shown on two-dimensional plots with 45-deg line (black dots). Red dots in lower panels denote their differences (Attend L – Attend R). For SD, CV, and median (**B-D**), only differences are shown as red dots as well as their mean ± SE. The IPI became shorter (**A** and **D**) and more regular (**B** and **C**) in the contra-attention hemisphere. These results were invariant whether we focused on IPIs in a delta-to-beta band (2 - 30 Hz) or an alpha-to-beta band (8 - 30 Hz). \*\**p* < 0.01, \*\*\**p* < 0.001.

### Statistical analysis

A purpose of the present study was to measure changes in oscillatory signals related to perception and attention. An effect of attention was investigated using the data in the cue period. For each of 102 sensor positions, data in Attend-Left condition (LL + LR) were compared with those in Attend-Right condition (RL + RR) by a paired *t*-test (N = 22 vs. 22). Resultant *t*-maps over a spatial layout of 102 sensor positions were shown in **Figures 2 and 3**. For example, attention-related suppression of alpha power (Worden et al., 2000; Wyart and Tallon-Baudry, 2009; Bauer et al., 2014) was shown in upper right panel in **Figure 2D**. The Attend-Right condition induced the suppression of alpha power in the contralateral left hemisphere (Attend-L > Attend-R, colored in red), while the Attend-Left condition induced alpha suppression in the right hemisphere (Attend-L < Attend-R, blue). A problem of multiple comparisons (caused by a repetition of *t*-test at 102 sensor positions) was resolved by controlling false discovery rate (FDR). For each *t*-map, a statistical threshold was adjusted for multiple testing with Benjamini-Hochberg correction (Benjamini and Hochberg, 1995) with the *q*-value set at < 0.05. Sensors with a significant difference after this correction were indicated by white rectangles. For means, SDs, and medians of IPIs, this correction of multiple comparisons was performed over all 102 sensor positions. For CV, the correction was made over a subset of sensors (sensors of interest or SOIs) showing a significant (*p* < 0.05) difference in SDs between Attend-Left vs. Attend-Right (**Fig. 3B**). We defined those SD-based SOIs to ensure that an attention-related decrease in CV (see **Results**) reflected a decrease in SD (numerator of CV), not an increase in mean IPI (denominator of CV), at the same sensor position.

We then investigated changes in powers and IPIs related to visual perception (**Fig. 6** and **Fig. 7**). Data in Stimulus-Left condition (LL + RL) were compared with Stimulus-Right condition (LR + RR). Other procedures (sensor-space *t*-map and correction of multiple comparisons, etc.) were the same as those on attention-related changes.

## Results

### Behavioral data

Hit rates for a target stimulus and false-alarm rates for non-target stimuli were 95.0 ± 2.2 % and 0.28 ± 0.11 %, respectively (mean ± SE across subjects). Reactions times measured from an onset of a grating were 665.9 ± 23.6 ms in LFV and 665.7 ± 22.2 ms in RVF. No difference in reaction times was observed between LVF and RVF (*t*(21) = 0.02, *p* = 0.98, Cohen’s *d* = 0.002).

### Oscillatory powers and IPIs in the cue period

**Figure 2A** shows TF (time-frequency) spectra around the cue period (−1200 to 0 ms) when participants directed their attention into LVF (an average of LL and RL trials, expressed as “LL+RL” hereafter). To obtain an overview of all 102 sensor positions, we classified those into 26 areas (25 areas of four sensor positions plus one area of two sensor positions). Spectra at four (or two) positions within each area were averaged together (areal-mean analyses (Suzuki et al., 2014)). Consistent with previous studies (Worden et al., 2000; Wyart and Tallon-Baudry, 2009; Bauer et al., 2014), an allocation of attention induced continuous suppression of alpha power (8-12 Hz) in the contralateral hemisphere (**Fig. 2B and C**). Statistical comparisons of powers between Attend-Left (LL + LR) and Attend-Right (RL + RR) conditions (sensor-space *t*-maps, **Fig. 2D**) revealed attention-related ERD in the contralateral hemisphere not only in alpha rhythm but also in delta (2-4 Hz), theta (5-7 Hz), and beta (13-30 Hz) rhythms.

Based on those results in power analysis, we computed distributions of IPIs with a band pass-filter of 2-30 Hz (delta-to-beta band), comparing their mean and variance (SD) between Attend-Left and Attend-Right conditions. A sensor-space *t*-map of mean IPI (Attend L vs. R) is shown in the left panel of **Figure 3A**. Attending to LVF reduced the mean IPI length over the right visual cortex (Attend L < R, colored in blue), while attending to RVF reduced the IPIs over the left visual cortex (Attend L > R, colored in red). The same results were obtained when we analyzed IPIs of alpha-to-beta rhythms using a band-pass filter of 8-30 Hz (middle panel of **Fig. 3A**). Individual data (N = 22) in alpha-to-beta IPIs at two representative sensors are shown in two-dimensional plots in the right panels. In the left hemisphere (Sensor 1), most black dots were located below a diagonal (45-deg) line, indicating that IPIs in Attend-Right condition were shorter than those in Attend-Left condition. Differences of mean IPI (Attend L - R) were thus positive in most participants (red dots). These patterns were reversed in the right hemisphere (Sensor 2) in which mean IPI was shorter in Attend-Left than Attend-Right conditions.

Attention-related changes in SD of IPIs are shown in **Figure 3B**. We found a significant decrease in SD in the contra-attention hemisphere, indicating that neural oscillations in delta-to-beta band (left panel) and alpha-to-beta band (middle panel) became more periodic by an allocation of attention to a contralateral VF. Regularity of oscillatory signals was further evaluated by CV (coefficient of variation), which was the SD divided by the mean. Significant decreases in the contra-attention hemisphere (**Fig. 3C**, middle) showed that attention not only induced the alpha/beta ERD but also enhanced the regularity of neural rhythms in the same regions.

One possible reason for the attention-related decrease in mean IPI (**Fig. 3A**) is an inhibition of transient events by ERD. It is widely known that high-power oscillation in a given frequency (e.g. 14 Hz) can emerge intermittently in neural waveforms (e.g. spindles and beta burst) (Shin et al., 2017). Such transient events might produce IPIs far longer than average (outlier IPIs). The mean IPI in the contra-attention visual cortex became shorter (**Fig. 3A**) because the alpha/beta ERD in this region (**Fig. 2D**) might inhibit an emergence of those transient bursts, thereby eliminating outlier IPIs. To examine this possibility, we computed a median of an IPI distribution, a statistical measure less influenced by outliers. Results revealed an attention-related decrease in the contralateral hemisphere (**Fig. 3D**), which suggests that the shortening of IPIs took place as a whole (not resulting from the elimination of outliers). Our results thus cannot be explained by the inhibition of transient bursts by alpha/beta ERD.

Supplementary analyses showed that those results were robust and not influenced by settings of the band-pass filter in the IPI analysis. **Figure 4** displays the results when we used a finite impulse response (FIR) filter, instead of an infinite impulse response (IIR) filter, to identify IPIs. Significant reductions of IPI measures were observed in the contra-attention hemisphere. In **Figure 5**, we compared results when an order of the IIR filter (Butterworth, zero-phase) was set at 2, 3, and 4. Attention-related reductions in alpha-to-beta IPIs were observed in all orders, indicating that our results cannot be explained by any contributions of frequency components outside the pass-band (e.g. theta and gamma waves).

**Figure 4.**
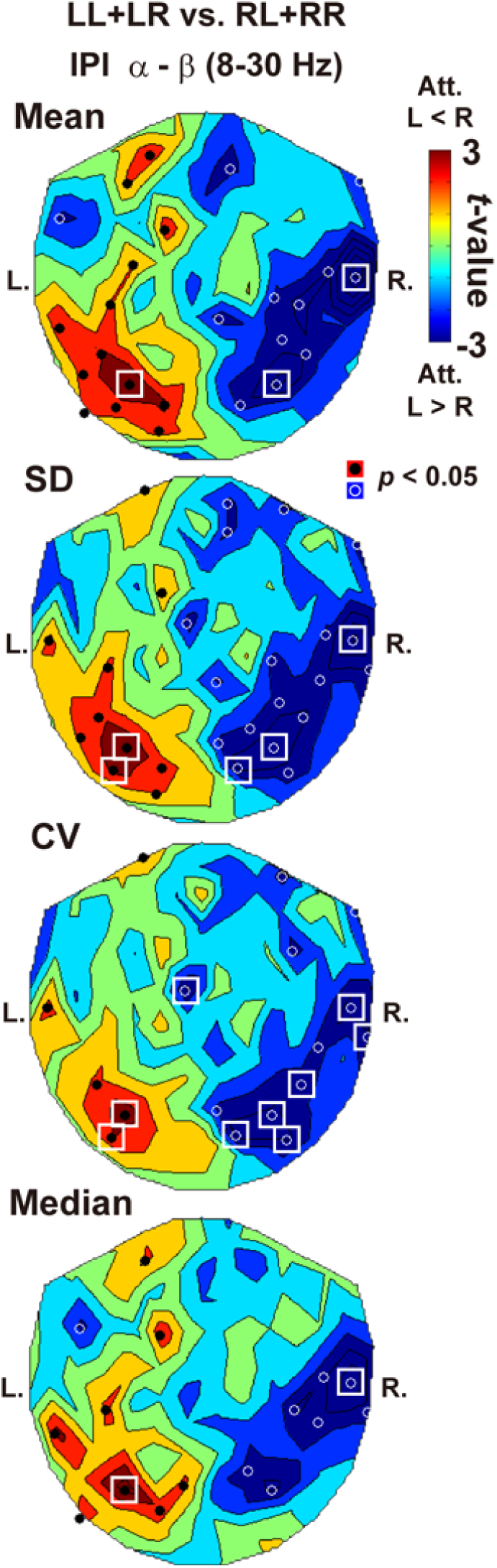
Results of the IPI analysis with an FIR (finite impulse response) filter. Although IPIs were measured using an IIR (infinite impulse response) filter (Butterworth, zero-phase) in **Figure 3**, here we show results of the IPI analysis in the cue period using an FIR filter (order: 1053). As in **Figure 3**, significant reductions of IPIs were observed in the contra-attention hemisphere. Our results therefore did not depend on a type (FIR or IIR) of digital filter.

**Figure 5.**
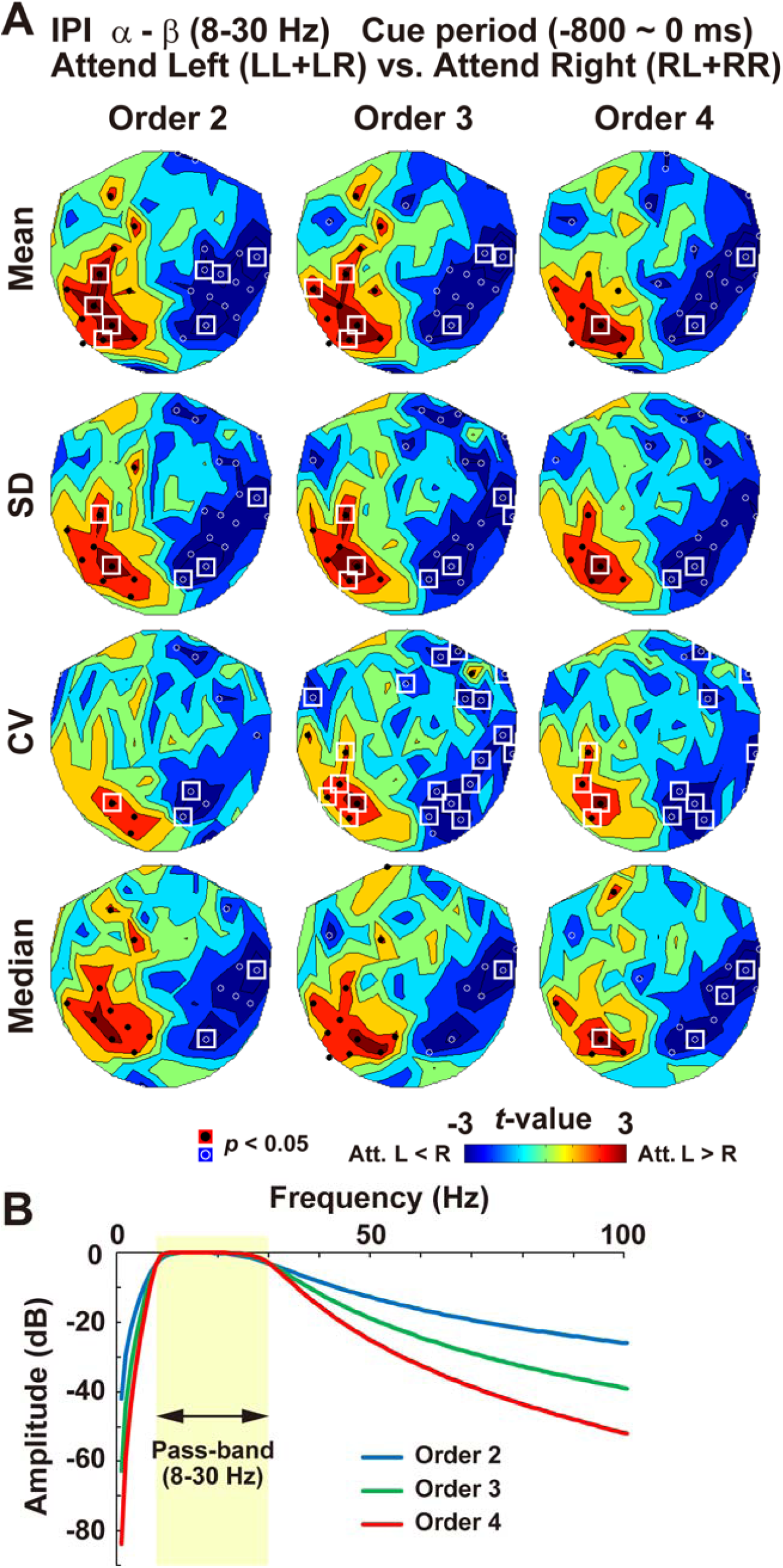
An effect of filter order. Although we used the IIR filter with an order of 3 in **Figure 3**, here we compared results the IPI analysis when the order was set at 2, 3, and 4 (**A**). Frequency responses in each filter (pass-band: 8-30 Hz) are also shown in panel **B**. Attention-related reductions in alpha-to-beta IPIs were observed in all orders. Those results indicate that our results cannot be explained by any contributions of frequency components outside the pass-band, such as theta and gamma waves (if our results had resulted from the components outside the pass-band, an effect of those components should have been smaller along with an increase in filter order).

### Oscillatory powers and IPIs in the target period

We then analyzed the data in the target period. Presenting a task stimulus induced alpha/beta ERD starting from 200 ms (**Fig. 6A**), consistent with previous studies (Minami et al., 2014; Kulashekhar et al., 2016). After 500 ms, this ERD was replaced by an increase in power (ERS), which reflected an inhibitory control of button-press movements to a non-target grating (see below). The ERD from 200 to 500 ms was more clearly seen in the contra-than ipsi-stimulus hemispheres (**Fig. 6B-D**). Since sensor-space *t*-maps of powers between Stimulus-Left (LL + RL) and Stimulus-Right (LR + RR) conditions indicated the contralateral ERD in alpha and beta band (**Fig. 7A**), we investigated a distribution of IPIs at 8-30 Hz (**Fig. 7B**). Results reveled that IPIs became faster (mean and median) and more regular (SD and CV) in the contra-stimulus hemisphere.

**Figure 6.**
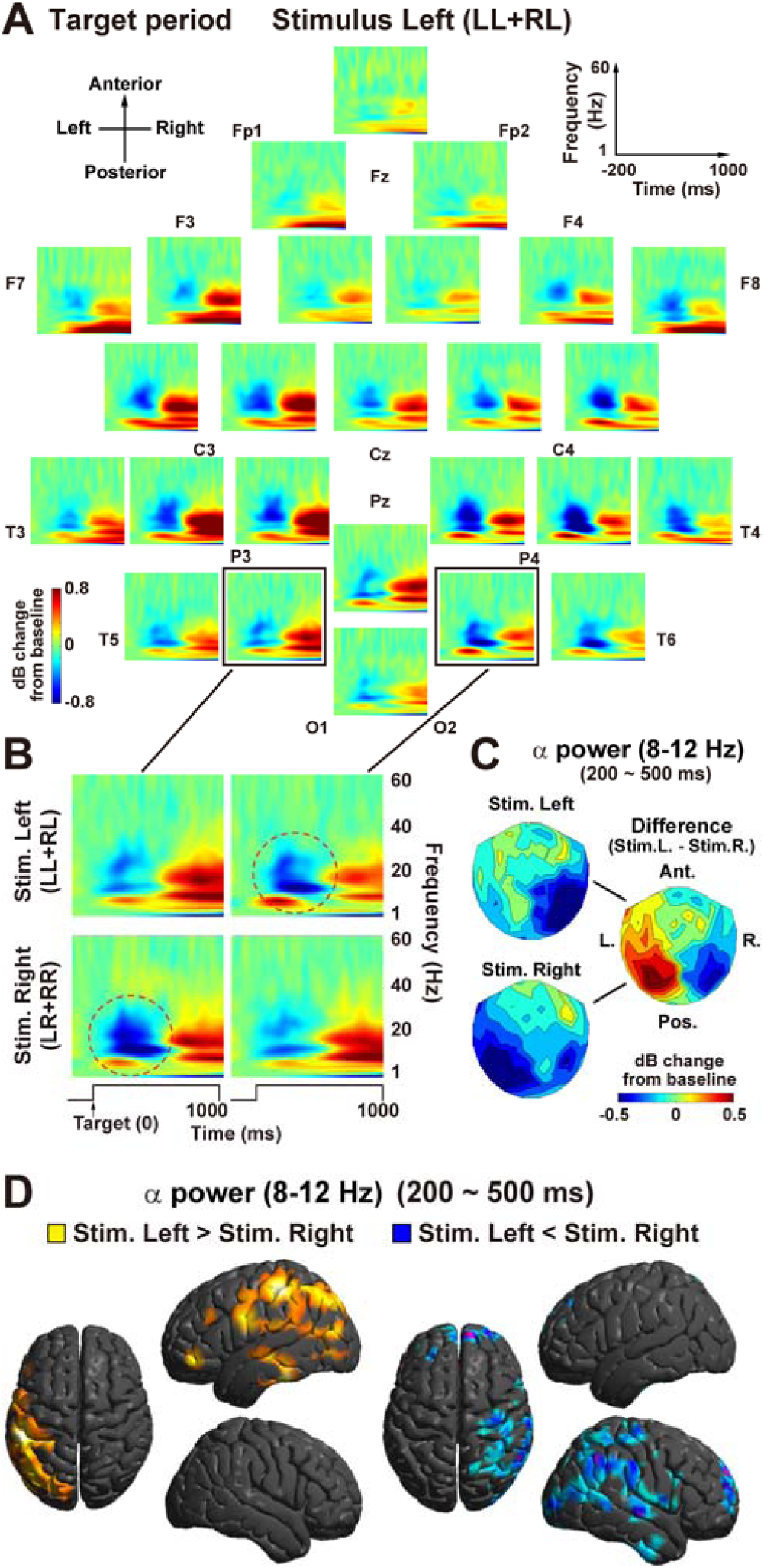
Results of power analysis in the target period. (**A**) Areal-mean spectra when a grating was presented at LVF (Stimulus-Left conditions). (**B**) Enlarged TF spectra over the bilateral visual regions. Presenting a stimulus predominantly suppressed alpha power over the contralateral visual region (dotted red circles). (**C**) Contour maps of alpha power over 102 sensor positions. Mean power in a time window of 200 - 500 ms are compared between Stimulus-Left and Stimulus-Right conditions. A differential map (Stimulus-Left minus Stimulus-Right) was characterized by an increase and decrease in alpha power in the left and right visual regions, respectively. (**D**) Anatomical source locations of the alpha suppressions estimated by SPM12. Regions with decreased alpha power in Stimulus-Left conditions are shown in blue, while those in Stimulus-Right conditions are shown in yellow.

**Figure 7.**
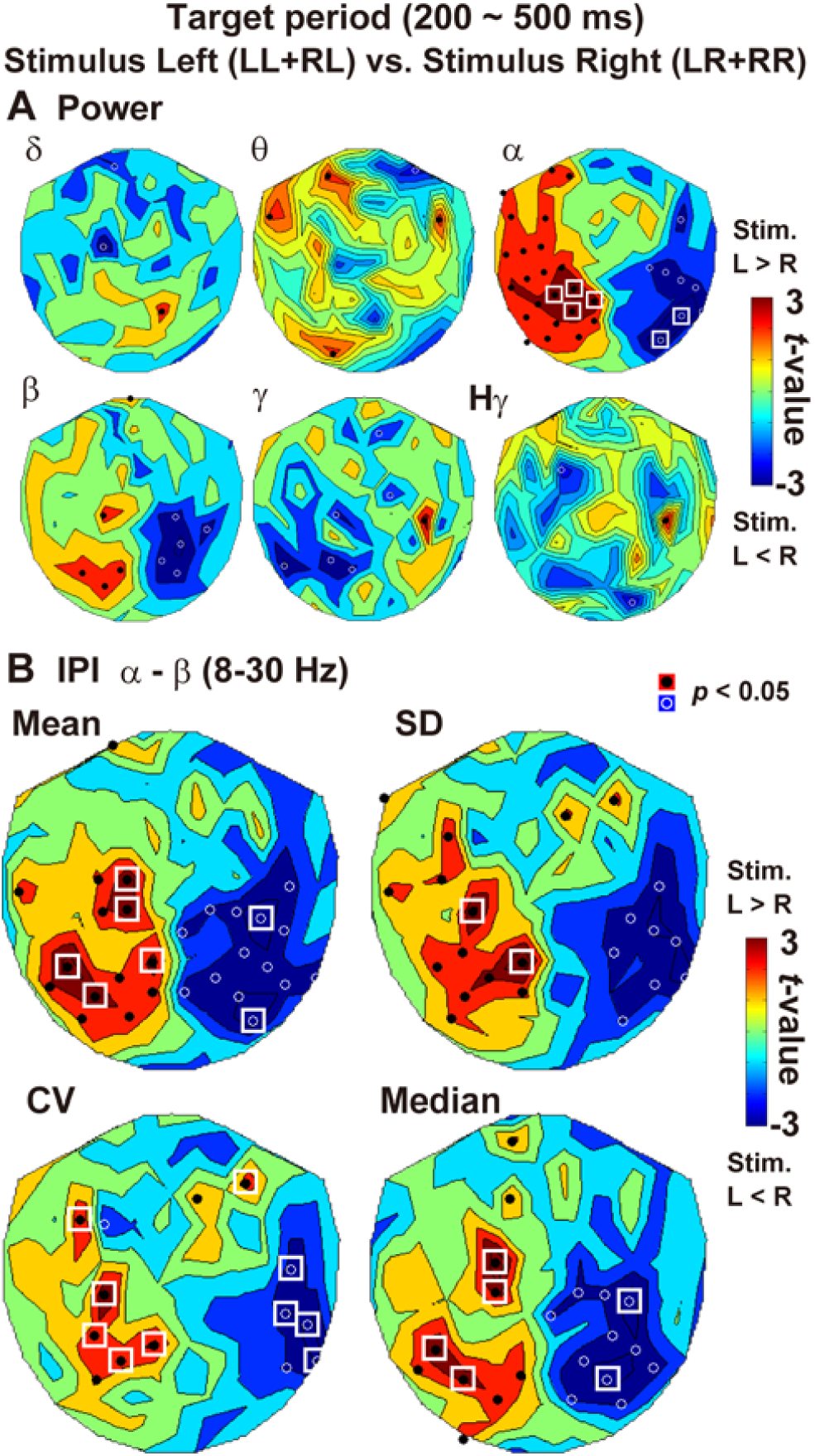
Changes in powers and IPIs in the target period (200 – 500 ms). (**A**) Statistical *t*-maps of the power analysis (Stimulus-Left vs. Stimulus-Right). Consistent with previous studies, presenting a visual stimulus induced suppression in alpha and beta bands in the contra-stimulus hemisphere. (**B**) The *t*-maps of the IPI analysis. Since the stimulus-induced ERD was observed in alpha and beta bands (**A**), a band-pass filter to measure IPIs was set at 8-30 Hz. The IPI became shorter (mean and median) and more regular (SD and CV) in the contra-stimulus hemisphere.

### NoGo inhibition and an increase in IPIs

Previous studies reported an increase in beta power in the frontal (motor-related) cortex during successful stop trials in Go/NoGo tasks (Jenkinson and Brown, 2011). A recent study further investigated a relationship between an inhibitory control and top-down attention using the spatial-cuing Go/NoGo task (Hong et al., 2017). They found that, while a NoGo stimulus presented in an attended visual filed evoked various EEG components related to response inhibition (NoGo-N2 and NoGo-P3), those components were greatly reduced or absent when the NoGo stimulus was presented in an unattended visual field. These previous data indicate that, in the present study, more inhibitory control was necessary when non-target (NoGo) grating appeared in an attended hemifield (LL and RR trials) than when it appeared in an unattended hemifield (LR and RL trials).

Consistent with this prediction, our data in a late target period (500 - 1000 ms) showed prominent alpha/beta ERS in congruent (LL + RR) compared to incongruent (LR + RL) conditions (**Fig. 8A**). Importantly, the ERS was mainly seen in anterior regions of the left hemisphere, which was contralateral to button-press movements (right hand). Results of IPI analysis at 8-30 Hz are displayed in **Figure 8B**. We found a significant increase in mean and median IPIs in congruent (inhibitory) compared to incongruent (non-inhibitory) conditions, although no change was observed in SD and CV. Those results showed that neural rhythms of alpha-to-beta band were slowed down in the frontal region when inhibitory control was necessary.

**Figure 8.**
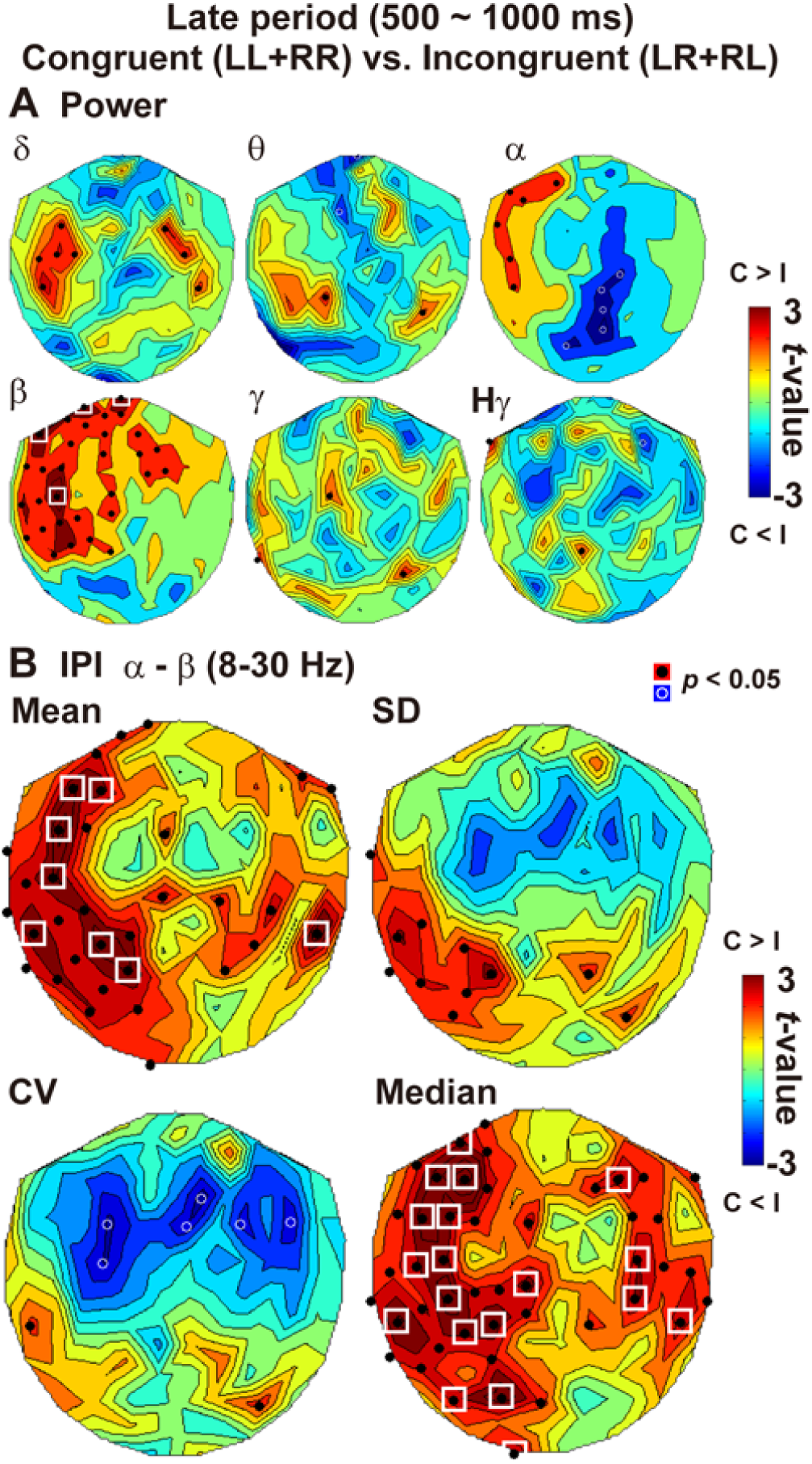
Changes in powers and IPIs in the late target period (500 - 1000 ms). (**A**) The *t*-maps of the power analysis. In congruent trials, a non-target grating was presented at an attended visual field (LL and RR), while it was shown in an unattended field in incongruent trials (LR and RL). No-go (non-target) stimulus at the attended hemifield induced neural activity related for response inhibition (Hong et al., 2017), as was shown by an increase in beta power (congruent > incongruent, lower left panel) in the frontal cortex (Jenkinson and Brown, 2011) contralateral to button-press movements (right hand). (**B**) The *t*-maps of the IPI analysis. An inhibitory control of button-press movements in the congruent trial was associated with an increase in mean and median IPIs. No change was observed in SDs in the left frontal regions. Those results suggest that the human brain achieved inhibitory control to a non-target stimulus by slowing down the neural rhythm in the left frontal (motor-related) regions without affecting its regularity.

### A leading role of beta rhythm in IPI modulations

We have observed IPI changes in alpha-to-beta band related to attention (**Fig. 3**), visual perception (**Fig. 7B**), and response inhibition (**Fig. 8B**). Which band (alpha or beta) played a more prominent role in those IPI modulations? We computed mean IPI separately for alpha (8-12 Hz), beta (13-30 Hz) as well as gamma (31-59 Hz) and high-gamma (60-100 Hz) bands, showing *t*-maps in **Figure 9A**. Differences between critical conditions (e.g. Attend L vs. Attend R in the left panels) were most clearly seen in beta band in all three functions (attention, perception, and inhibitory control). Those results indicated a leading role of beta rhythms in IPI modulations (distributions of beta IPIs in a representative subject are shown in **Fig. 9B**).

**Figure 9.**
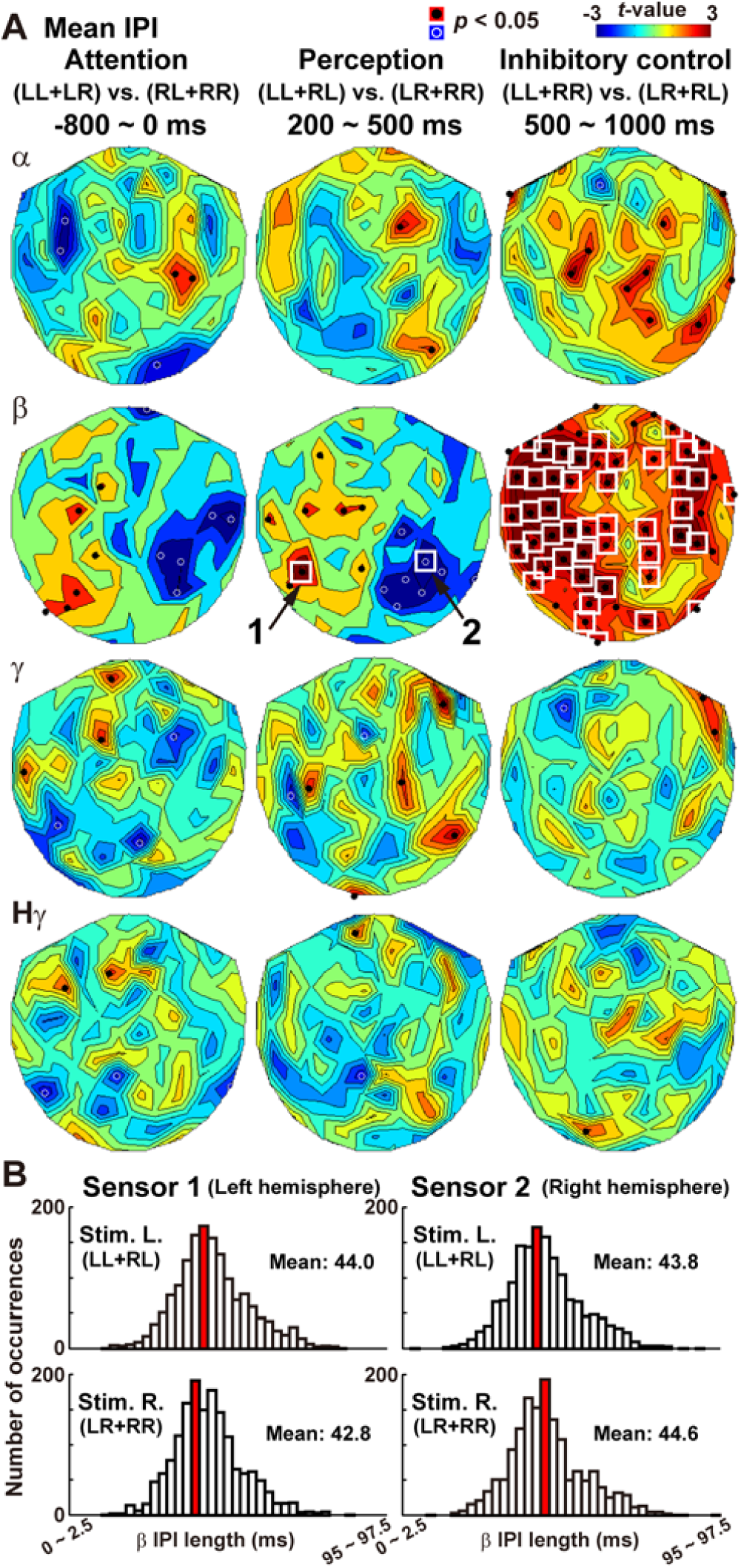
A leading role of beta rhythm in IPI modulations. (**A**) The *t*-maps of mean IPI separately calculated for alpha, beta, gamma, and high-gamma bands. Modulation of IPIs related to attention (Attend-Left vs. Attend-Right), perception (Stimulus-Left vs. Stimulus-Right), and inhibitory control (congruent vs. incongruent) are shown in left, middle, and right columns, respectively. Data in delta and theta bands are not shown because of small numbers of IPIs in those bands (especially in the target period of 200-500 ms). Note distinct changes of beta IPIs observed in all three functions (attention, perception, and inhibitory control). (**B**) Distributions of the beta IPIs in a representative subject. A red bar in each histogram denotes a bin (width: 2.5 ms) with a maximum counts (mode). Presenting a grating in RVF shortened the beta IPIs over the left visual cortex (Sensor 1), whereas a grating in LVF shortened those over the right visual cortex (Sensor 2).

### Dissociation of the power and IPI analyses

So far, we have reported a tight coupling between oscillatory powers and IPIs; alpha/beta ERD in the contra-attention and contra-stimulus hemisphere was associated with shortened IPIs (**Fig. 3 and 7**), while alpha/beta ERS in the left frontal region was linked with an extended IPIs (**Fig. 8**). In a final section of **Results**, we would report a dissociation of those two measures.

It is well known that a stimulus presented in an attended location evokes stronger neural responses in the sensory cortex than a stimulus presented in an unattended location (Hillyard and Anllo-Vento, 1998), a phenomenon called the attentional modulation (AM). Interestingly, a recent study investigating the stimulus-induced beta suppression reported that the AM was more prominent in the *ipsi-stimulus* than contra-stimulus hemisphere (van Ede et al., 2014). We successfully replicated their findings in **Figure 10A**. A task stimulus in LVF produced stronger beta suppression when subjects had attended to LVF (LL trials) than when they had attended to RVF (RL trials), which corresponds to the AM (left panel of **Fig. 10A**). Numbers of sensors showing the significant (*p* < 0.05) AM was 27 in the left hemisphere and 4 in the right hemisphere. A chi-square test yielded a significant bias (*χ*^2^(1) = 25.20, *p* = 0.0000005, *φ*= 0.512), indicating that the AM was more prominent in ipsi-stimulus (left) hemisphere. A comparison between RR and LR trials also showed the AM more distinct in ipsi-stimulus (right) hemisphere (right panel of **Fig. 10A**, *χ*^2^(1) = 9.19, *p* = 0.002, *φ* = 0.309).

**Figure 10.**
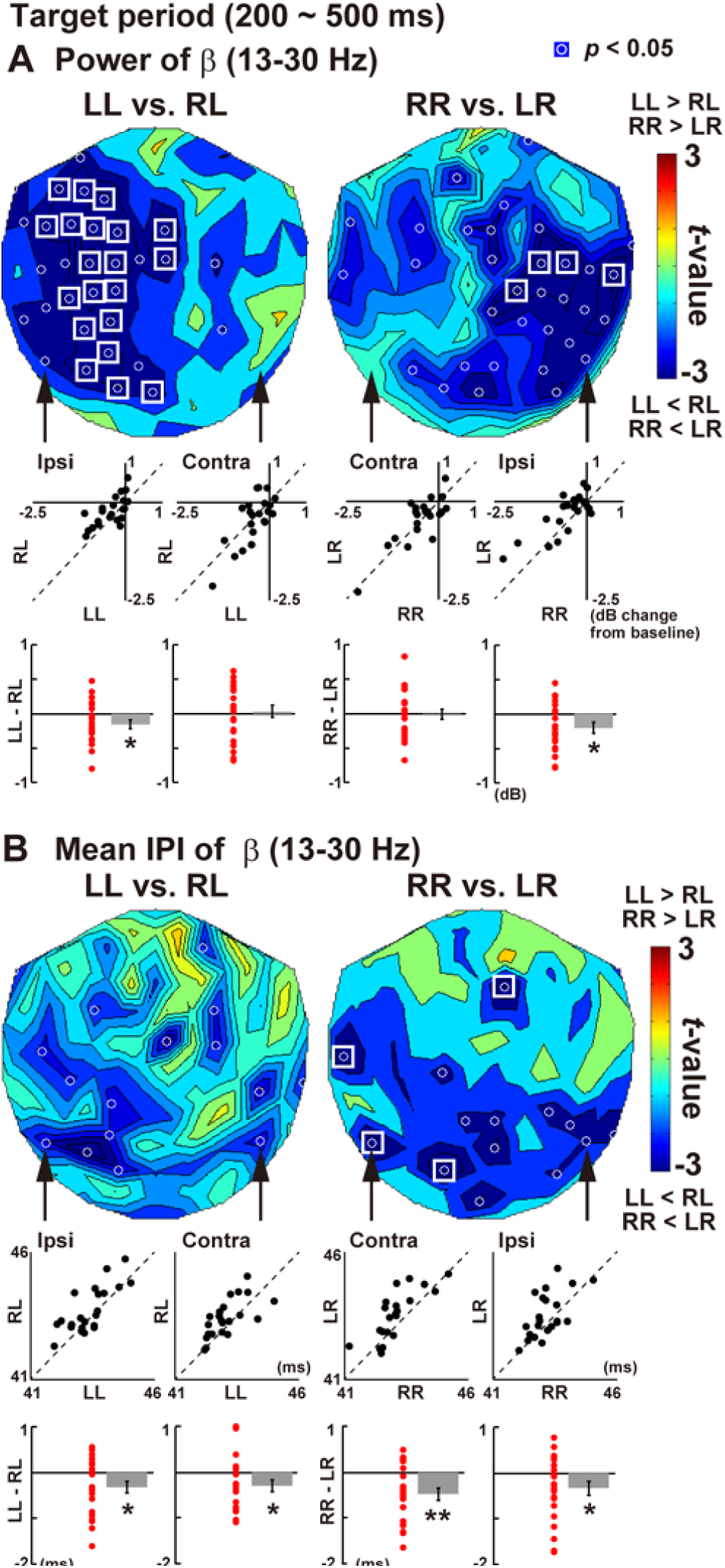
Dissociation between the power and IPI analyses. (**A**) The *t*-map of beta power at the target period (congruent vs. incongruent trials). Consistent with a previous study (van Ede et al., 2014), an effect of congruency (attentional modulation) was prominent in the ipsi-stimulus (rather than contra-stimulus) hemisphere. For example, when a grating was presented in LVF (left panel), significant differences in beta suppression between congruent (LL) and incongruent (RL) trials were observed in the left (ipsi-stimulus) hemisphere. Individual data (mean beta power at 200-500 ms) at sensors indicated by arrows are shown as black dots in the two dimensional plots (e.g. LL vs. RL). Individual differences between congruent and incongruent trials are also shown as the red dots as well as their mean ± SE. Note that, although magnitudes of beta suppression were greater in contra-stimulus than ipsi-stimulus hemispheres, an effect of congruency (e.g. LL vs. RL) did not reach significance in the contra-stimulus hemisphere (because of strong beta suppression in both congruent and incongruent trials). (**B**) The *t*-map of mean IPI of beta rhythm (congruent vs. incongruent trials). An effect of congruency was observed bilaterally (not lateralized to the ipsi-stimulus hemisphere). Results of the IPI analyses therefore can dissociate from those of the power analysis; significant changes in IPIs can emerge from sensors with no change in powers.

This dominance of the ipsi-stimulus hemisphere on the AM (van Ede et al., 2014), however, was not observed in results of IPI analysis. As shown in **Figure 10B**, the AM in IPIs was bilateral. Although IPIs in congruent (LL and RR) condition were shorter than those incongruent (LR and RL) trials, this reduction of IPIs (AM) was equally observed in contra-stimulus and ipsi-stimulus hemispheres (LL vs. RL: *χ*^2^(1) = 0.30, *p* = 0.58, *φ* = 0.056, RR vs. LR: *χ*^2^(1) = 0.80, *p* = 0.37, *φ* = 0.091). Of particular interest were sensors in the contra-stimulus hemisphere that showed a significant AM of beta IPIs (**Fig. 10B**) despite no changes in beta power (**Fig. 10A**). Results of the IPI analysis thus can dissociate from those of the power analysis, suggesting separate neural mechanisms for controlling IPIs and powers.

## Discussion

In contrast to the prediction in **Introduction**, we presently found that alpha/beta ERD in task-relevant regions was associated with an increased regularity of oscillatory signals. Although regularity (periodicity) of neural oscillation has been measured using autocorrelation analysis (Red’ka and Mayorov, 2015), spectral entropy (Poza et al., 2012; Gomez-Pilar et al., 2016), sample entropy (Gomez and Hornero, 2010), auto-mutual information analysis (Gomez et al., 2007), Q factor (Lemercier et al., 2017), and lagged coherence (Fransen et al., 2015), no study has reported a perception-or attention-related increase in regularity in sensory areas (even an attention-related decrease in regularity was reported (Fransen et al., 2015)). We recorded neuromagnetic signals from the human brain at a high sampling rate of 4 kHz, directly depicting a distribution of IPIs. Results revealed that alpha/beta ERD was coupled to reductions in the mean and SD of their IPIs. Those reductions cannot be attributed to a general increase in vigilance or arousal level because they selectively took place in the contralateral hemisphere (bilateral changes would have been observed if they had reflected arousal). Significant reductions were also observed in the median of IPIs, which indicates that our results cannot be explained by an inhibition of transient neural events, e.g. the beta burst (Shin et al., 2017), by ERD. In contrast to a previous view associating the desynchronization with uncorrelated behaviors of neural population (Pfurtscheller and Lopes da Silva, 1999; Beaman et al., 2017), our results suggested a new role of the ERD to accelerate and regulate brain rhythms, probably in order to facilitate the neural processing in task-relevant regions.

### Dynamic modulation of IPIs linked to cognitive functions

It is widely accepted that electric and magnetic waveforms recorded from the brain are composed of multiple bands with different frequencies, each of which plays separate roles in mental functions. An oscillation in theta band is tightly coupled with memory functions (Knyazev, 2007; Buzsaki and Moser, 2013), while that in gamma band is thought to be important in attention, etc. (Engel et al., 2001; Fries, 2009). Based on this principle, many previous studies first decomposed EEG/MEG waveforms into 5 or 6 bands (e.g. delta, theta, alpha… etc.) and then calculated a “mean power” of each band by pooling data across neighboring frequencies (e.g. 2-4 Hz for theta and 30-60 Hz for gamma). In other words, subtle changes in frequency *within* each band have received little attention so far. In contrast to this static view, we presently found a subtle but dynamic shift of frequency in oscillation signals, as was shown by the reduction in mean IPI. For example, attending to RVF reduced a mean length of alpha-to-beta IPIs in the left visual cortex from 49.4 to 48.8 ms (**Fig. 3A**). This corresponds to a shift in dominant frequency from 20.24 to 20.49 Hz, which would be undetectable by the previous approach. Our data suggest a mechanism in the human brain that dynamically adjusts its oscillation in a sub-millisecond scale, depending on sensory inputs and an allocation of attention.

In addition to mean IPI, our data showed significant changes in SD and CV. Those reductions indicate that basic rhythms in the brain became faster (mean) and more regular (SD) during visual stimulation and attentional allocation. One intriguing idea is that those oscillation signals represent the rhythm of a clock for the neural computation (information processing) in the brain. The reduced mean and SD thus suggest that our brain might perceive a stimulus and control attention by regulating and accelerating the neural clock in task relevant regions. Although many theories and models have been proposed about neural mechanisms of perceptual consciousness (Dehaene et al., 2014; Tononi et al., 2016) and attention (Bisley, 2011; Mueller et al., 2017), the present data of IPIs would provide useful information to refine these models from a perspective of macro-level temporal dynamics of neural activity.

### Leading role of beta rhythm in IPI changes

For detailed analyses of IPIs, we depicted *t*-maps for each of traditional frequency bands (**Fig. 9A**). Results revealed a leading role of beta rhythm in modulating IPIs. This indicates that changes in IPIs are *not* a direct consequence of changes in oscillation amplitude. As shown by the power analysis, alpha wave showed the distinct ERD related to perception (LL+RL vs. LR+RR, **Fig. 7A**) and attention (LL+LR vs. RL+RR, **Fig. 2D**). Nevertheless, those decreases in alpha power elicited no changes in alpha IPIs in the same regions (**Fig. 9A**), indicating that a change in oscillation amplitude did not necessarily lead to a change in IPIs. In contrast, the perception- and attention-related ERD in beta rhythm was significant but less distinct than alpha band (**Fig. 2D** and **Fig. 7A**). Those modest changes in beta power, however, were accompanied by significant changes in beta IPIs (**Fig. 9A**). In brief, alpha rhythm played a leading role in modulating the amplitude of oscillations, while beta rhythm played a leading role in modulating IPIs. Compared to other frequency bands, functional roles of beta rhythm remain controversial and unclear (Engel and Fries, 2010; Spitzer and Haegens, 2017). Our results suggest a new and special role of beta oscillation to orchestrate brain rhythms in a sub-millisecond scale, depending on a presence/absence of visual stimuli and a spatial allocation (left/right) of attention.

On the other hand, the present data also showed that beta oscillation was necessary but not sufficient to fully control brain rhythms. The reductions in IPIs were more robustly observed when we investigated MEG waveforms comprising broader frequency components (2-30 Hz or 8-30Hz, **Fig. 3**) than when only the beta band was analyzed (13-30 Hz, **Fig. 9**). Although beta wave plays an important role in modulating IPIs, it achieved more stable control of brain rhythms by interacting with other bands of frequencies (e.g. alpha).

### Possibility and future directions of the IPI analysis

Our IPI analyses provided some data showing potentials of this method in future studies. In the later target period (500 - 1000 ms), for example, we observed a significant increase in mean IPI in the left frontal regions along with beta ERS (**Fig. 8**). This suggests that our brain made a “no-go” decision by slowing down neural rhythms in the left frontal (motor-related) cortex. The IPI analysis is thus applicable not only to an investigation of perception and attention but also to that of inhibitory control in the frontal cortex. Those results also showed a bi-directional relationship between beta power and beta IPIs; the ERD in beta band was linked to shorter IPIs (perception and attention), while the beta ERS was linked to longer IPIs (inhibitory control).

Another advantage of the IPI analysis was seen in results of the attention modulation (AM) (**Fig. 10**). Using the data at 200-500 ms, we compared oscillatory signals between congruent (LL and RR) and incongruent (LR and RL) trials. Power analyses (**Fig. 10A**) revealed stronger beta ERD to a grating in congruent than incongruent trials (AM), and this AM was more clearly seen in the ipsi-stimulus (rather than contra-stimulus) hemisphere (van Ede et al., 2014). On the other hand, the AM in IPI data was bilateral (**Fig. 10B**). Reduced IPIs in congruent compared to incongruent trials were observed not only in ipsi-stimulus but also in contra-stimulus hemispheres. Several sensors in the contra-stimulus hemisphere showed significant changes in beta IPIs despite a lack of change in beta power. These shortened IPIs in the contra-stimulus visual cortex might be directly related to shorter reaction times to a stimulus presented at an attended than unattended visual fields (Posner et al., 1980). Our IPI approach therefore would provide original data that cannot be detected through a traditional power analyses.

## Acknowledgments

This work was supported by KAKENHI Grants Number 22680022 and 26700011 from the Japan Society for the Promotion of Science (JSPS) for Young Scientists to Y.N. We thank Mr. Y. Takeshima (National Institute for Physiological Sciences, Japan) for his technical supports.

The authors declare no competing financial interest.

## References

Bauer M, Oostenveld R, Peeters M, Fries P (2006) Tactile spatial attention enhances gamma-band activity in somatosensory cortex and reduces low-frequency activity in parieto-occipital areas. J Neurosci 26:490–501.

Bauer M, Stenner MP, Friston KJ, Dolan RJ (2014) Attentional modulation of alpha/beta and gamma oscillations reflect functionally distinct processes. J Neurosci 34:16117–16125.

Beaman CB, Eagleman SL, Dragoi V (2017) Sensory coding accuracy and perceptual performance are improved during the desynchronized cortical state. Nat Commun 8:1308.

Benjamini Y, Hochberg Y (1995) Controlling the False Discovery Rate-a Practical and Powerful Approach to Multiple Testing. Journal of the Royal Statistical Society Series B-Methodological 57:289–300.

Berger H (1929) Uber das Elektrenkephalogramm des Menschen. Arch Psychiat Nervenkr 87:527–750.

Bisley JW (2011) The neural basis of visual attention. J Physiol 589:49–57.

Brainard DH (1997) The Psychophysics Toolbox. Spat Vis 10:433–436.

Buzsaki G, Moser EI (2013) Memory, navigation and theta rhythm in the hippocampal-entorhinal system. Nat Neurosci 16:130–138.

Cirelli LK, Bosnyak D, Manning FC, Spinelli C, Marie C, Fujioka T, Ghahremani A, Trainor LJ (2014) Beat-induced fluctuations in auditory cortical beta-band activity: using EEG to measure age-related changes. Front Psychol 5:742.

de Pesters A, Coon WG, Brunner P, Gunduz A, Ritaccio AL, Brunet NM, de Weerd P, Roberts MJ, Oostenveld R, Fries P, Schalk G (2016) Alpha power indexes task-related networks on large and small scales: A multimodal ECoG study in humans and a non-human primate. Neuroimage 134:122–131.

Dehaene S, Charles L, King JR, Marti S (2014) Toward a computational theory of conscious processing. Curr Opin Neurobiol 25:76–84.

Engel AK, Fries P (2010) Beta-band oscillations--signalling the status quo? Curr Opin Neurobiol 20:156–165.

Engel AK, Fries P, Singer W (2001) Dynamic predictions: oscillations and synchrony in top-down processing. Nat Rev Neurosci 2:704–716.

Fransen AM, Dimitriadis G, van Ede F, Maris E (2016) Distinct alpha- and beta-band rhythms over rat somatosensory cortex with similar properties as in humans. J Neurophysiol 115:3030–3044.

Fransen AM, van Ede F, Maris E (2015) Identifying neuronal oscillations using rhythmicity. Neuroimage 118:256–267.

Fries P (2005) A mechanism for cognitive dynamics: neuronal communication through neuronal coherence. Trends Cogn Sci 9:474–480.

Fries P (2009) Neuronal gamma-band synchronization as a fundamental process in cortical computation. Annu Rev Neurosci 32:209–224.

Fujioka T, Ross B, Trainor LJ (2015) Beta-Band Oscillations Represent Auditory Beat and Its Metrical Hierarchy in Perception and Imagery. J Neurosci 35:15187–15198.

Gomez-Pilar J, Martin-Santiago O, Suazo V, de Azua SR, Haidar MK, Gallardo R, Poza J, Hornero R, Molina V (2016) Association between EEG modulation, psychotic-like experiences and cognitive performance in the general population. Psychiatry Clin Neurosci.

Gomez C, Hornero R (2010) Entropy and Complexity Analyses in Alzheimer’s Disease: An MEG Study. Open Biomed Eng J 4:223–235.

Gomez C, Hornero R, Abasolo D, Fernandez A, Escudero J (2007) Analysis of the magnetoencephalogram background activity in Alzheimer’s disease patients with auto-mutual information. Comput Methods Programs Biomed 87:239–247.

Hillyard SA, Anllo-Vento L (1998) Event-related brain potentials in the study of visual selective attention. Proc Natl Acad Sci U S A 95:781–787.

Hong X, Wang Y, Sun J, Li C, Tong S (2017) Segregating Top-Down Selective Attention from Response Inhibition in a Spatial Cueing Go/NoGo Task: An ERP and Source Localization Study. Sci Rep 7:9662.

Jenkinson N, Brown P (2011) New insights into the relationship between dopamine, beta oscillations and motor function. Trends Neurosci 34:611–618.

Jensen O, Kaiser J, Lachaux JP (2007) Human gamma-frequency oscillations associated with attention and memory. Trends Neurosci 30:317–324.

Jensen O, Mazaheri A (2010) Shaping functional architecture by oscillatory alpha activity: gating by inhibition. Front Hum Neurosci 4:186.

Klimesch W (2012) alpha-band oscillations, attention, and controlled access to stored information. Trends Cogn Sci 16:606–617.

Knyazev GG (2007) Motivation, emotion, and their inhibitory control mirrored in brain oscillations. Neurosci Biobehav Rev 31:377–395.

Kulashekhar S, Pekkola J, Palva JM, Palva S (2016) The role of cortical beta oscillations in time estimation. Hum Brain Mapp 37:3262–3281.

Lemercier CE, Holman C, Gerevich Z (2017) Aberrant alpha and gamma oscillations ex vivo after single application of the NMDA receptor antagonist MK-801. Schizophr Res 188:118–124.

Minami T, Noritake Y, Nakauchi S (2014) Decreased beta-band activity is correlated with disambiguation of hidden figures. Neuropsychologia 56:9–16.

Mueller A, Hong DS, Shepard S, Moore T (2017) Linking ADHD to the Neural Circuitry of Attention. Trends Cogn Sci 21:474–488.

Nishitani N, Hari R (2002) Viewing lip forms: cortical dynamics. Neuron 36:1211–1220.

Noguchi Y, Kimijima S, Kakigi R (2015) Direct behavioral and neural evidence for an offset-triggered conscious perception. Cortex 65:159–172.

Osipova D, Takashima A, Oostenveld R, Fernandez G, Maris E, Jensen O (2006) Theta and gamma oscillations predict encoding and retrieval of declarative memory. J Neurosci 26:7523–7531.

Palva S, Palva JM (2011) Functional roles of alpha-band phase synchronization in local and large-scale cortical networks. Front Psychol 2:204.

Pelli DG (1997) The VideoToolbox software for visual psychophysics: transforming numbers into movies. Spat Vis 10:437–442.

Pfurtscheller G, Lopes da Silva FH (1999) Event-related EEG/MEG synchronization and desynchronization: basic principles. Clin Neurophysiol 110:1842–1857.

Posner MI, Snyder CR, Davidson BJ (1980) Attention and the detection of signals. J Exp Psychol 109:160–174.

Poza J, Gomez C, Bachiller A, Hornero R (2012) Spectral and Non-Linear Analyses of Spontaneous Magnetoencephalographic Activity in Alzheimer’s Disease. Journal of Healthcare Engineering 3:299–321.

ReD’ka IV, Mayorov OY (2015) Effect of Restriction of Visual Afferentation on the Rhythmic Organization of Alpha EEG Activity. Neurophysiology 47:225–233.

Shin H, Law R, Tsutsui S, Moore CI, Jones SR (2017) The rate of transient beta frequency events predicts behavior across tasks and species. Elife 6.

Spitzer B, Haegens S (2017) Beyond the Status Quo: A Role for Beta Oscillations in Endogenous Content (Re)Activation. eNeuro 4.

Suzuki M, Noguchi Y, Kakigi R (2014) Temporal dynamics of neural activity underlying unconscious processing of manipulable objects. Cortex 50:100–114.

Tadel F, Baillet S, Mosher JC, Pantazis D, Leahy RM (2011) Brainstorm: a user-friendly application for MEG/EEG analysis. Comput Intell Neurosci 2011:879716.

Taube JS (2010) Interspike interval analyses reveal irregular firing patterns at short, but not long, intervals in rat head direction cells. J Neurophysiol 104:1635–1648.

Tononi G, Boly M, Massimini M, Koch C (2016) Integrated information theory: from consciousness to its physical substrate. Nat Rev Neurosci 17:450–461.

van Ede F, de Lange FP, Maris E (2014) Anticipation increases tactile stimulus processing in the ipsilateral primary somatosensory cortex. Cereb Cortex 24:2562–2571.

van Ede F, Koster M, Maris E (2012) Beyond establishing involvement: quantifying the contribution of anticipatory alpha- and beta-band suppression to perceptual improvement with attention. J Neurophysiol 108:2352–2362.

Worden MS, Foxe JJ, Wang N, Simpson GV (2000) Anticipatory biasing of visuospatial attention indexed by retinotopically specific alpha-band electroencephalography increases over occipital cortex. J Neurosci 20:RC63.

Wyart V, Tallon-Baudry C (2009) How ongoing fluctuations in human visual cortex predict perceptual awareness: baseline shift versus decision bias. J Neurosci 29:8715–8725.

